# Effect of Moisture Content of Paddy (Phka Rumduol) on the Quality of White Rice (*Oryza sativa* L.)

**DOI:** 10.64898/2026.11.18.767487

**Authors:** In Sokra

## Abstract

This study evaluated the impact of paddy moisture content on the quality parameters of white rice produced from Cambodia’s premium fragrant variety, Phka Rumduol (*Oryza sativa* L.). A completely randomized design (CRD) was employed to evaluate the effect of moisture content on rice quality. Paddy samples were adjusted to four target moisture levels (10%, 12%, 14%, and 16%) and subsequently milled under controlled conditions. The CRD ensured unbiased allocation of samples across all treatments, minimizing experimental error during both moisture conditioning and milling processes. The results showed that moisture content had a significant effect (p < 0.05) on all measured quality parameters. Paddy at 14% moisture content produced the best results, with a head rice yield of 63.46 ± 3.00%, and a broken rice percentage of only 4.86 ± 0.65%. Grain appearance was also well preserved, with rice bran yield of 10.67 ± 0.42%, rice husk yield of 18.52 ± 2.72%, and yellow kernels averaging just 0.79 ± 0.02%. In contrast, moisture levels below 12% increased kernel brittleness, leading to breakage rates as high as 26.28 ± 0.92%, while levels above 16% caused grain deformation and discoloration. These findings highlight the importance of maintaining paddy moisture at 14% prior to milling to optimize recovery and preserve the high quality of Phka Rumduol rice. The study provides practical guidance for Cambodian rice processors aiming to improve product quality and strengthen competitiveness in international markets.

## 1. Introduction

Rice (*Oryza sativa* L.) is one of the major cereal crops produced worldwide, serving as a primary food source for more than 50% of the global population, particularly in Asia (Mohapatra & Sahu, 2022). The quality of milled white rice is essential for consumer preference, market value, and economic returns for producers. Among the various factors affecting paddy quality, moisture content before milling plays a key role in determining grain integrity and overall product quality (Tong et al., 2019). Proper postharvest moisture management is crucial to minimize breakage, maximize whole kernel yield, and preserve quality parameters. In Cambodia, paddy rice is not only a staple food but also a key agricultural export, with fragrant varieties such as Phka Rumduol, Somali, and Phka Malis being particularly valued (Thong et al., 2021). Phka Rumduol, a long-grain aromatic rice, is internationally recognized for its pleasant fragrance, soft texture, and elongation upon cooking. However, its premium quality can be compromised if postharvest handling—particularly drying and moisture control—is not properly managed. The moisture content of paddy rice before milling significantly impacts the mechanical resistance of grains during dehusking and polishing. Insufficient drying leads to high moisture content, making kernels susceptible to deformation and breakage, while excessive drying increases brittleness, resulting in higher fractions of broken rice (Bunna et al., 2018). Therefore, identifying the optimal moisture content for milling Phka Rumduol is critical for maintaining its market competitiveness. Previous studies on rice milling quality have reported varying optimal moisture levels depending on rice variety, milling technology, and environmental conditions. For instance, research on Jasmine rice in Thailand suggested that a moisture content of 13–15% before milling maximizes head rice yield (Chitsuthipakorn & Thanapornpoonpong, 2021).

Similarly, studies in Cambodia demonstrated that 14–15% moisture content is ideal for minimizing breakage in local varieties (Bunna et al., 2018; Sopheap et al., 2020). However, other research has reported different optimal moisture levels: Firouzi and Alizadeh (2013) found that Iranian rice varieties performed best at 8–9% moisture, while Siebenmorgen et al. (2007) reported that higher moisture levels (18–23.5%) improved milling yields in certain U.S. long-grain rice varieties. These discrepancies suggest that rice varieties respond differently to moisture levels, necessitating variety-specific studies to determine optimal postharvest practices. Additionally, moisture content affects other rice quality parameters including whiteness, yellowness, and aroma retention. High moisture content in paddy rice promotes microbial growth and discoloration, leading to off-color grains that reduce market value (Shafiekhani et al., 2018; Somaly et al., 2025). Given that Phka Rumduol is typically exported to premium markets, maintaining its visual and sensory qualities is essential for sustaining Cambodia’s reputation as a producer of high-quality rice.

Despite the economic importance of Phka Rumduol, there is limited research on the specific moisture requirements for optimizing its milling quality, leaving a gap in knowledge regarding Cambodia’s flagship fragrant rice. This study aims to determine the effect of paddy moisture content (ranging from 10% to 16%) on the milling quality of Phka Rumduol, with a focus on whole kernel yield, head rice yield, broken rice percentage, and yellow kernel incidence. The findings will provide practical recommendations for Cambodian rice farmers and millers to optimize moisture content for enhanced milling recovery and product quality. This research supports the broader goal of strengthening Cambodia’s rice export industry and ensuring the sustainability of its premium rice market.

## 2. Materials and Methods

### 2.1 Raw Materials

Phka Rumduol paddy rice (*Oryza sativa* L.), a premium aromatic rice variety in Cambodia, was used in this study. The paddy was collected immediately after harvest from a local farm in Thmor Kol Commune, Battambang Province, Cambodia. Four moisture content treatments were prepared: 10%, 12%, 14%, and 16% (wet basis). Each treatment consisted of three replicates, with each replicate containing 1 kg of paddy rice, resulting in a total of 12 kg used in the experiment. All samples were manually cleaned to remove dirt, stones, and other foreign materials prior to moisture adjustment.

### 2.2 Experimental Design

The experiment followed a completely randomized design (CRD) to investigate the effect of different paddy moisture contents on the quality of white rice. Four moisture content treatments were established: 10%, 12%, 14%, and 16% (wet basis). Each treatment included three replicates, and each replicate consisted of 1 kg of paddy rice, totaling 12 experimental units. After moisture adjustment, all samples were processed under identical conditions, including dehulling and polishing. Quality assessments focused on three key parameters: whole kernel percentage, head rice yield, and yellow kernel percentage.

- The treatments were designated as follows:
- T1: 10% moisture content of paddy
- T2: 12% moisture content of paddy
- T3: 14% moisture content of paddy
- T4: 16% moisture content of paddy

### 2.3 Process of Experiment

All paddy rice was harvested from the same field in Thmor Korl commune with an initial moisture sontent of approximately 23%. The freshly harvested rice was immediately subjected to drying using a temperature-controlled mechanical dryer, operating between 30°C and 40°C, until the desired moisture levels were achieved. Four target moisture contents were prepared: 10%, 12%, 14%, and 16% (wet basis). Once the specified moisture contents were reached, all samples were milled using the same milling machine. For each treatment, 1 kg of paddy rice was used. Milling was conducted at a low speed to minimize grain breakage and ensure uniform quality. The milled rice from each treatment was then evaluated for quality parameters, including whole kernel percentage, head rice yield, and yellow kernel percentage.

### 2.4 Determination of Head Rice Percentage

To determine the head rice percentage, 1000 g of milled white rice from each replicate was taken and evenly spread on a black-and-white board. Using tweezers, head rice grains—defined as kernels longer than 75% of the whole grain length—were carefully separated and weighed. The percentage of head rice was calculated following the method described by Siebenmorgen et al. (2013).

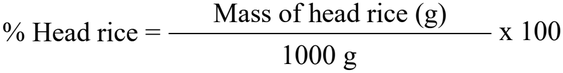

### 2.5 Determination of Broken Rice Percentage

To determine the broken rice percentage, 1000 g of milled white rice from each replicate was taken and evenly spread on a black-and-white board. Using tweezers, the broken rice grains—defined as rice kernels that are less than 75% of the full grain length—were carefully separated and weighed. The percentage of broken rice was calculated following the method escribed by Aghayeghazvini et al. (2009).

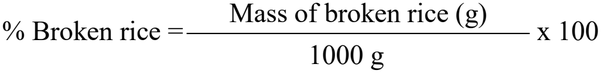

### 2.6 Determination of Rice Bran Percentage

To determine the rice bran percentage, 1000 g of milled white rice from each replicate was evenly spread on a black-and-white board. The bran layer was carefully separated using tweezers and then weighed. The percentage of rice bran was calculated following the method described by Siebenmorgen et al. (2013).

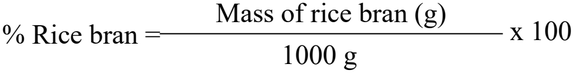

### 2.7 Determination of Rice Husk Percentage

To determine the rice husk percentage, 1000 g of milled white rice from each replicate was evenly spread on a black-and-white board. The bran layer was carefully separated using tweezers and then weighed. The percentage of rice husk was calculated following the method described by Siebenmorgen et al. (2013).

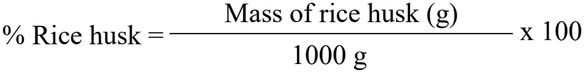

### 2.8 Determination of Yellow Kernel

To determine the yellow kernel percentage, 1000 g of milled white rice from each replicate was evenly spread on a clean, flat surface. Yellow kernels—identified by their distinct discoloration compared to normal white rice grains—were visually separated and collected using tweezers. The percentage of yellow kernel was calculated following the method described by Wang et al. (2002).

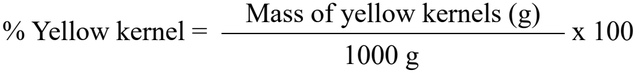

### 2.9 Statistical Analysis

All experiments were conducted in triplicate, and the data were expressed as mean ± standard deviation (SD). Statistical analysis was performed using GraphPad Prism software (version 10.4.2, Windows, San Diego, California, USA; www.graphpad.com). One-way analysis of variance (ANOVA) was used to assess significant differences among treatment means. When ANOVA indicated significance (p < 0.05), Tukey’s Honestly Significant Difference (HSD) test was applied for multiple comparisons.

## 3. Results

### 3.1 Changes of Head Rice

**Fig. 1** presents the mean head rice percentages obtained from different moisture content treatments. The whole kernel percentage varied significantly with moisture levels (p < 0.01). Treatment T3 (14% moisture content) produced the highest head rice percentage (63.46 ± 3.00%), which was significantly greater than all other treatments. T2 (12%) showed a moderate value (49.52 ± 2.08%), followed by T4 (16%) at 42.85 ± 2.03%. The lowest percentage was observed in T1 (10%), with a value of 37.48 ± 1.90%. These findings suggest that a moisture content of 14% is optimal for maximizing whole kernel recovery during milling, whereas both lower and higher moisture levels result in increased kernel breakage.

**Fig. 1.**
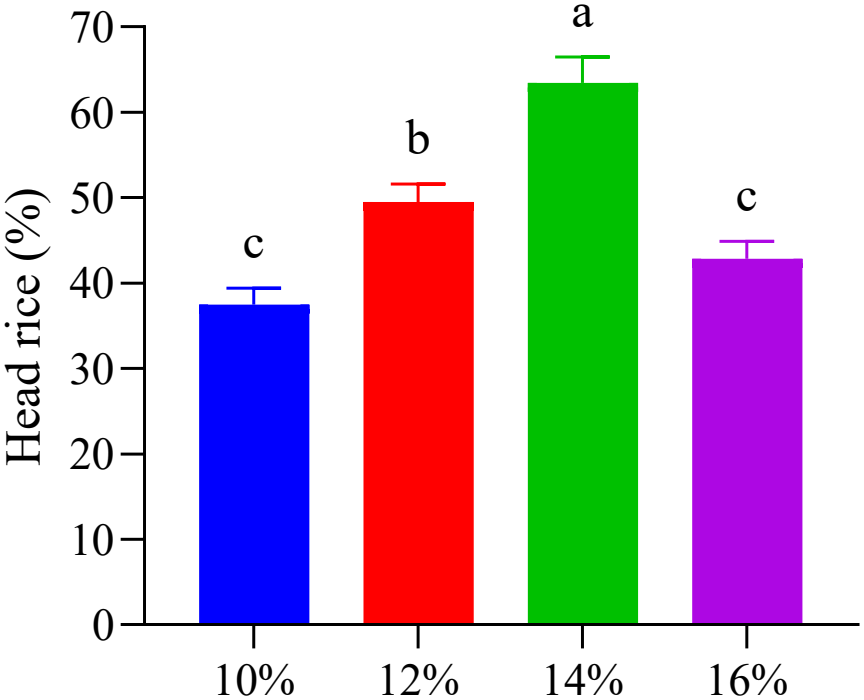
Effect of different moisture contents (10%, 12%, 14%, and 16%) on the percentage of head rice in milled rice. Bars represent mean ± standard deviation (n = 3). Different letters above bars indicate statistically significant differences at p < 0.05 according to Tukey’s HSD test.

### 3.2 Changes in Broken Rice

**Fig.2** indicates the mean percentages of broken rice obtained from samples with different moisture contents during milling. Moisture content significantly affected the broken rice percentage (p < 0.01). The lowest percentage of broken rice was observed in T3 (14% moisture content), with 4.86 ± 0.65%, which was significantly lower than all other treatments. T2 (12%) also showed a relatively low value of 15.14 ± 0.55%. In contrast, higher percentages were recorded in T1 (10%) and T4 (16%), with 26.28 ± 0.92% and 21.37 ± 1.43%, respectively. These results suggest that a moisture content of 14% minimizes rice breakage during milling, while both lower and higher moisture levels result in greater grain damage.

**Fig. 2.**
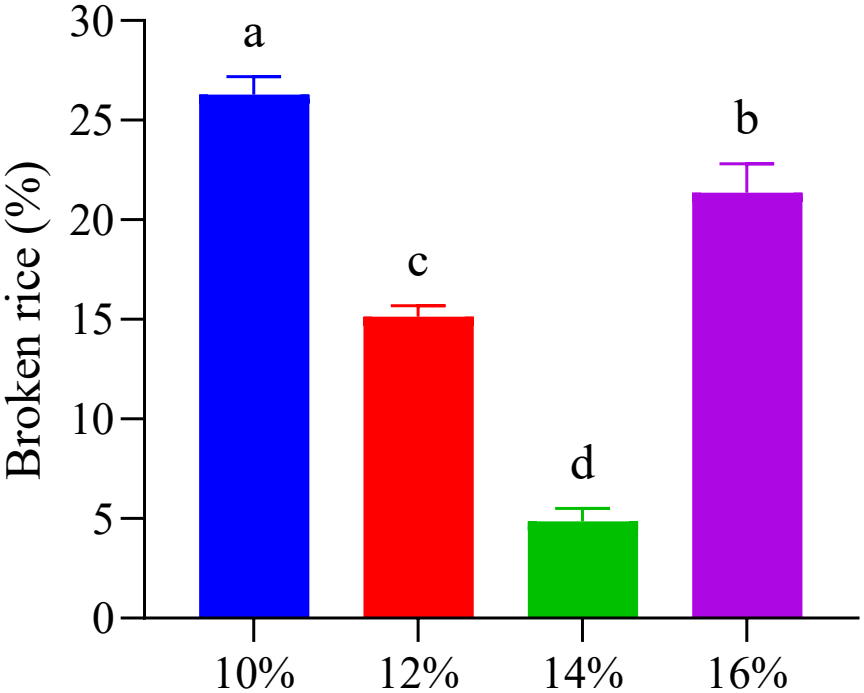
Effect of different moisture contents (10%, 12%, 14%, and 16%) on the percentage of broken rice after storage. Bars represent mean ± standard deviation (n = 3). Different letters above bars indicate statistically significant differences at p < 0.05 according to Tukey’s HSD test.

### 3.3 Changes in Rice Bran

**Fig.3** illustrates the mean percentages of rice bran observed in milled rice at different moisture contents. The rice bran percentage did not vary significantly among treatments (p > 0.05). The values ranged narrowly from 11.43 ± 0.90% in T4 (16% moisture content) to 11.22 ± 1.38% in T1 (10%), with T2 (12%) and T3 (14%) showing intermediate values of 10.67 ± 0.61% and 10.67 ± 0.42%, respectively. These results suggest that moisture content in the tested range had minimal impact on bran formation during milling.

**Fig. 3.**
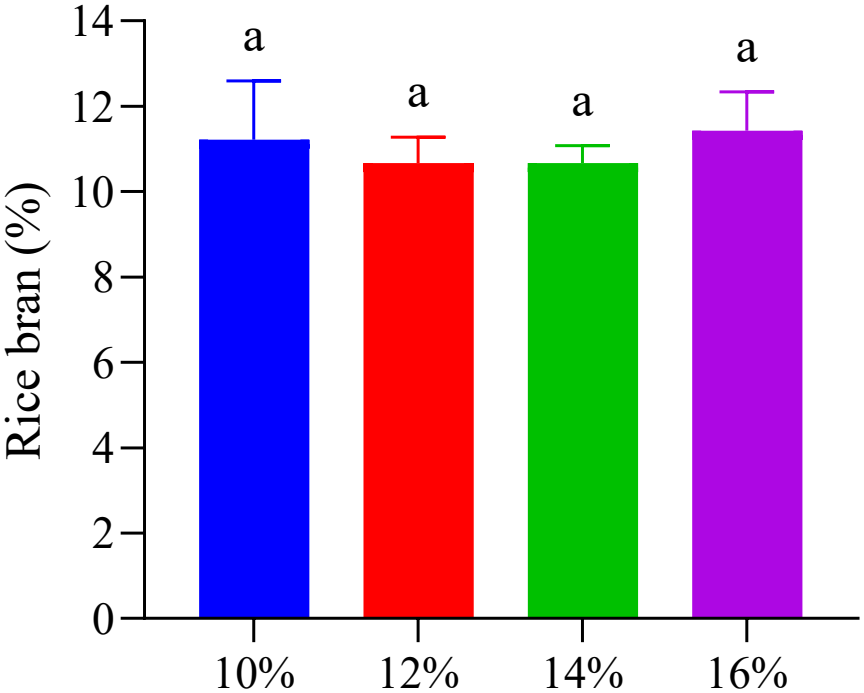
Effect of different moisture contents (10%, 12%, 14%, and 16%) on the percentage of rice bran after milling. Bars represent mean ± standard deviation (n = 3). Different letters above bars indicate statistically significant differences at p < 0.05 according to Tukey’s HSD test.

### 3.4 Changes in Rice Husk

**Fig.4** shows the average rice husk percentages in milled rice samples subjected to varying moisture content treatments. Statistical analysis indicated no significant differences among the treatments (p > 0.05). Rice husk content remained relatively consistent across all moisture levels, with the highest value recorded in T1 (10%) at 21.03 ± 0.67% and the lowest in T3 (14%) at 18.53 ± 2.72%. T2 (12%) and T4 (16%) produced intermediate values of 20.57 ± 0.21% and 20.96 ± 1.67%, respectively. Overall, the findings imply that the tested moisture content range had little influence on husk retention during the milling process.

**Fig. 4.**
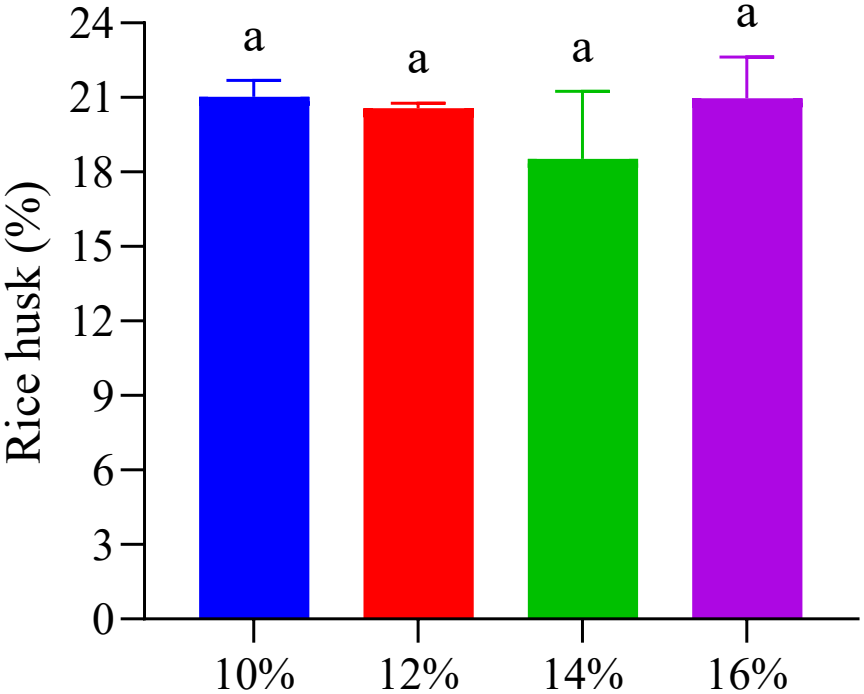
Effect of different moisture contents (10%, 12%, 14%, and 16%) on the percentage of rice husk in milled rice. Bars represent mean ± standard deviation (n = 3). Different letters above bars indicate statistically significant differences at p < 0.05 according to Tukey’s HSD test.

#### 3.5 Changes in Yellow Kernel

**Fig.5** illustrates the average percentage of yellow kernels in milled rice across different moisture content treatments. The analysis revealed no statistically significant variation among treatments (p > 0.05). Yellow kernel percentages remained fairly uniform, ranging from 0.79 ± 0.02% in T3 (14% moisture content) to 0.82 ± 0.01% in T2 (12%). Treatments T1 (10%) and T4 (16%) yielded comparable values of 0.80 ± 0.02% and 0.80 ± 0.01%, respectively. These results suggest that within the tested moisture range, moisture content had minimal effect on the occurrence of yellow kernels during the milling process.

**Fig. 5.**
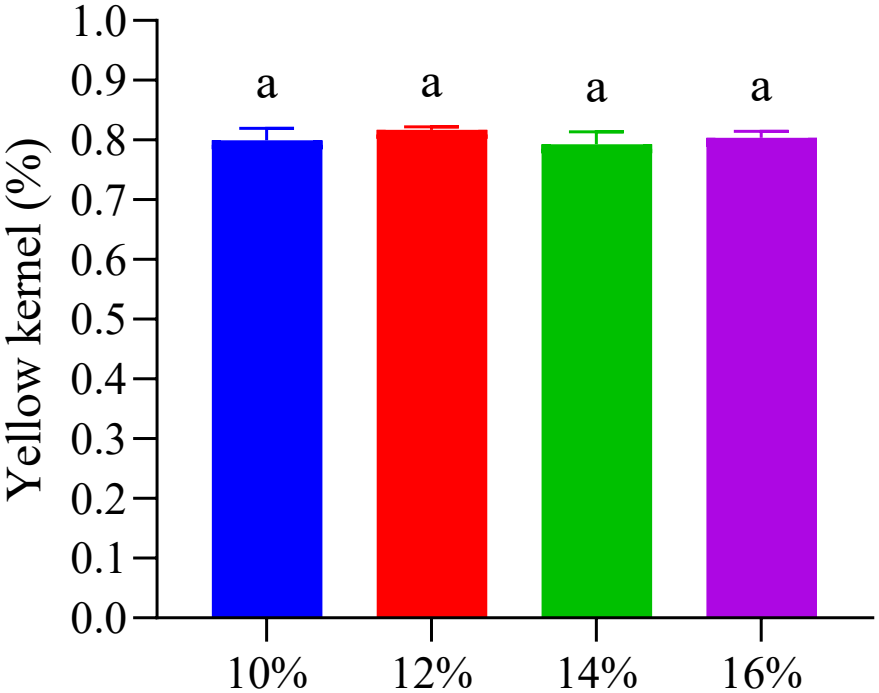
Effect of different moisture contents (10%, 12%, 14%, and 16%) on the percentage of yellow kernel in milled rice. Bars represent mean ± standard deviation (n = 3). Different letters above bars indicate statistically significant differences at p < 0.05 according to Tukey’s HSD test.

The milling yields of rice fractions were significantly affected by the grain’s moisture content before milling (10%, 12%, 14%, and 16%). Head rice yield peaked at 14% moisture (634.6 ± 30.04 g), followed by 12% moisture (495.2 ± 20.81 g), while 10% and 16% moisture resulted in significantly lower yields (374.4 ± 19.00 g and 428.47 ± 20.30 g, respectively (p < 0.01). In contrast, broken rice yield was highest at 10% moisture (262.77 ± 9.17 g) and lowest at 14% moisture (48.63 ± 6.47 g), with intermediate values at 16% (213.73 ± 14.32 g) and 12% moisture (151.37 ± 5.51 g; p < 0.01). No significant differences were observed in rice bran yield (p > 0.05), rice husk yield (p > 0.05), or yellow kernel incidence (p > 0.05) across moisture levels, indicating these parameters were unaffected by pre-milling moisture variation (**Table 1**).

**Table 1.** Effect of different moisture contents on whole kernel weight, head rice weight, broken rice weight, and yellow kernel content in milled rice. Values are presented as mean ± SD (n = 3). Different letters in the same row indicate significant differences at p < 0.05 according to Tukey’s HSD test.

| Parameter (g) | T1 (10%) | T2 (12%) | T3 (14%) | T4 (16%) | p-value |
|---|---|---|---|---|---|
| Head rice | 374.4 ± 19.00<sup>c</sup> | 495.2 ± 20.81<sup>b</sup> | 634.6 ± 30.04<sup>a</sup> | 428.47 ± 20.30<sup>c</sup> | 0.001 |
| Broken rice | 262.77 ± 9.17<sup>a</sup> | 151.37 ± 5.51<sup>c</sup> | 48.63 ± 6.47<sup>d</sup> | 213.73 ± 14.32<sup>b</sup> | 0.001 |
| Rice bran | 112.17 ± 13.82<sup>a</sup> | 106.67 ± 6.11<sup>a</sup> | 106.67 ± 4.16<sup>a</sup> | 114.33 ± 9.02<sup>a</sup> | 0.66 |
| Rice husk | 210.33 ± 6.66<sup>a</sup> | 205.67 ± 2.08<sup>a</sup> | 185.33 ± 27.15<sup>a</sup> | 209.60 ± 16.69<sup>a</sup> | 0.27 |
| Yellow kernel | 8.00 ± 0.20<sup>a</sup> | 8.17 ± 0.06<sup>a</sup> | 7.93 ± 0.21<sup>a</sup> | 8.03 ± 0.12<sup>a</sup> | 0.38 |

Superscript letters (a, b, c) indicate significant differences among treatments within each row.

## 4. Discussion

The results of whole kernel recovery suggest that a moisture content of 14% is optimal for maximizing whole kernel yield during milling. Both lower and higher moisture contents lead to increased kernel breakage, highlighting the critical role of precise moisture control in rice postharvest processing. This finding aligns with Alim et al. (2024), who reported that drying paddy rice to moisture levels that are either too high or too low reduces whole kernel yield. Proper moisture content enhances grain hardness and elasticity, making kernels less prone to cracking and breakage during dehusking and polishing. The significance of moisture content for the milling quality of Phka Rumduol, a premium Cambodian rice variety, also depends on additional quality traits such as grain size and structural integrity, as noted by Sopheap et al. (2020). However, contrasting findings have been reported in other studies. For example, Firouzi and Alizadeh (2013) found that in Iran, paddy rice with a moisture content between 8–9% yielded the highest proportion of whole kernels. Similarly, Kim & Lee (2013) reported that the ideal moisture content for producing high-quality white rice falls between 7% and 15%. These discrepancies may be attributed to regional differences in rice varieties, milling equipment, and climatic conditions. Furthermore, Khomkaew (2022) supported the present findings, demonstrating that a moisture content of 14% prior to milling is most effective in maximizing yield. This study also found significant differences in head rice yield among treatments, with 14% moisture content producing the highest proportion. This aligns with Bunna et al. (2018), who reported that in Cambodia, the optimal moisture content for milling to achieve high head rice yield ranges from 14% to 15%. However, not all studies agree. Siebenmorgen et al. (2007) found that increasing moisture content to 18.7–23.5% was optimal for maximizing head rice yield, suggesting that moisture’s effect may depend on how rice kernels respond mechanically under different postharvest treatments. Kruszelnicka et al. (2022) also indicated that increasing moisture content alters the mechanical behavior of grains, which can improve milling outcomes under certain conditions. Nevertheless, our study confirms that exceeding the optimal moisture range does not always improve head rice yield in Phka Rumduol. Müller et al. (2022) supported this conclusion, finding that drying paddy rice to 14–15% moisture content was optimal for this variety to achieve high head rice yield. Regarding broken rice, the results clearly showed that 14% moisture content produced the lowest proportion of broken kernels, while both lower and higher moisture levels increased breakage. This supports the work of Thompson and Mutters (2006), who concluded that maintaining paddy rice at 14% moisture content before milling effectively reduces broken rice. Dilday (1989) also found that the percentage of broken kernels increased significantly as grain moisture content at harvest decreased. Excessively dry grains become brittle and prone to fracture, whereas overly moist grains are more susceptible to bran and husk resistance and stress fractures during processing. This study resulted of bran and husk in average of 30% which is consistent to Phuwadlpaisarn (2021) revealed that rice husk and rice bran from cultivar grown in different location of Thailand resulted around 30% in average. Additionally, moisture content did not significantly affect the yellow kernel rate in Phka Rumduol. This finding is consistent with Khema et al. (2022), who reported that yellow kernels in white rice—a key quality parameter for Cambodian rice—are primarily influenced by storage conditions rather than moisture content within the 10–18% range. Yellowing in paddy rice depends on multiple factors during storage, including moisture content, microbial growth, environmental conditions, fungal activity, and processing methods (Dillahunty et al., 2001; Sokra et al., 2024).

## 5. Conclusion

This study demonstrates that paddy moisture content is a critical factor in optimizing the milling quality of Cambodia’s premium fragrant rice, Phka Rumduol. The results confirm that 14% moisture content is the optimal level, yielding the highest whole kernel and head rice recovery while minimizing broken grains and discoloration. Deviations from this level compromise quality: below 12%, kernels become brittle and prone to breakage, while above 16%, excessive moisture leads to deformation and yellowing. These findings highlight the necessity of strict moisture control before milling. To maximize quality, efficiency, and market value, a target moisture content of 14% should be adopted—ensuring Cambodian fragrant rice maintains its competitive edge in domestic and global markets.

## Acknowledgment

This research received no specific grant from any funding agency in the public, commercial, or not-for-profit sectors. The author acknowledges the technical support and platform resources provided by Research Foundation (www.researchfoundation.top). All findings and contents presented in this article remain the sole responsibility of the author.

## Notes

### Competing Interest Statement

The authors have declared no competing interest.

